# The relationship between dynamic functional network connectivity and spatial orientation in healthy young adults

**DOI:** 10.1101/2021.02.14.431143

**Authors:** Mohammad S. E. Sendi, Charles A. Ellis, Robyn L. Miller, David H. Salat, Vince D. Calhoun

## Abstract

Spatial orientation is essential to interacting with a physical environment, and better understanding it could contribute to a better understanding of a variety of diseases and disorders that are characterized by deficits in spatial orientation. Many previous studies have focused on the relationship between spatial orientation and individual brain regions, though in recent years studies have begun to examine spatial orientation from a network perspective. This study analyzes dynamic functional network connectivity (dFNC) values extracted from over 800 resting-state fMRI recordings of healthy young adults (age 22-37 years) and applies unsupervised machine learning methods to identify neural brain states that occur across all subjects. We estimated the occupancy rate (OCR) for each subject, which was proportional to the amount of time that they spent in each state, and investigated the link between the OCR and spatial orientation and the state-specific FNC values and spatial orientation controlling for age and sex. Our findings showed that the amount of time subjects spent in a state characterized by increased connectivity within and between visual, auditory, and sensorimotor networks and within the default mode network while at rest corresponded to their performance on tests of spatial orientation. We also found that increased sensorimotor network connectivity in two of the identified states negatively correlated with decreased spatial orientation, further highlighting the relationship between the sensorimotor network and spatial orientation. This study provides insight into how the temporal properties of the functional brain connectivity within and between key brain networks may influence spatial orientation.

## INTRODUCTION

Spatial orientation is the capacity to understand where one’s body is in relation to the external physical environment [1], which enables one to execute actions that physically interact with that environment. Spatial orientation is essential to our ability to function in a physical world. A better understanding of the neural mechanisms associated with spatial orientation in healthy individuals could eventually provide a better understanding of the mechanisms by which deficits in spatial orientation are produced in traumatic brain injuries [2], vascular dementia [3], Alzheimer’s disease [4], developmental topographical disorientation [5], and other disorders. Traditional studies of spatial orientation typically focus on its relationship with individual brain regions. However, in recent years, analyses have begun to explore the links between spatial orientation and brain network functional connectivity [6,7]. While some studies have begun to analyze how brain networks relate to spatial orientation, they have not considered how resting-state brain network dynamics relate to spatial orientation skills. There is good reason to think that dynamic functional network connectivity (dFNC) might provide insight into spatial orientation.

Dynamic FNC has been applied to identify network states associated with cognitive functions like processing speed, fluid intelligence, and working memory [8], as well as neurological disorders including Alzheimer’s disease and mental disorders such as schizophrenia, depression, autism, and ADHD [9–12]. As such, we propose the use of dFNC to gain insight into the relationship between resting-state brain network dynamics and spatial orientation. Because dFNC involves the analysis of FNC at multiple time points, it enables the identification and evaluation of network state dynamics that correspond to various neurological functions and is uniquely suited for the analysis of resting-state data. Moreover, it has the potential to provide novel insights into our current understanding of the neural mechanisms associated with spatial orientation. Utilizing the dynamical property of dFNC, we analyze resting-state fMRI (rs-fMRI) recordings from more than 800 healthy young adults, identify several distinct network states, and identify a relationship between those states and performance on a test of spatial orientation skills.

## MATERIAL AND METHODS

This study analyzed rs-fMRI data from over 800 healthy young adults to identify states of dFNC that corresponded to performance on a test of spatial orientation skills.

### 2.1. Description of data

The dataset used for this study was composed of rs-fMRI data and scores from spatial orientation tasks collected from healthy young adults and made publicly available as part of the Human Connectome Project (HCP) [13]. The fMRI, demographic information, and cognitive scores are available for download on the HCP website (https://www.humanconnectome.org). This study was approved by the institutional review board of the Washington University - University of Minnesota Consortium of the Human Connectome Project (WU-Minn HCP).

#### 2.1.1. Participants

In the current study, we used rs-fMRI 0f 833 subjects (average age: 28.65; range: 22-37 years; female/male: 443/390) and their associated demographic and cognitive scores. The rs-fMRI data were collected on a Siemens Skyra 3T with a 32-channel RF receiver head coil. High-resolution T2*-weighted functional images were acquired using a gradient-echo EPI sequence with TE = 33.1 ms, TR= 0.72 s, flip angle = 52°, slice thickness = 2 mm, 72 s slices and 2 mm isotropic voxel, the field of view: 208×180 mm (RO×PE), and duration: 14:33 (min: sec).

#### 2.1.2 Spatial orientation test

The variable short Penn line orientation test (VSPLOT) was used to assess each subject’s spatial orientation processing [14]. In this test, two color-coded lines – a fixed line and a moveable line – with different orientations were presented to the subject via a screen. The subject had to click buttons to incrementally rotate the moveable line either clockwise or counterclockwise until it was parallel with the fixed line. Across trials, the relative location of the lines on the screen and the length of the moveable line varied, while the distance between the centers of the two lines and the length of the fixed line remained constant. Overall, there were 24 trials for each subject. The total number of the correct trials (VSPLOT_TC), median reaction time divided by the expected number of clicks to correctly align the lines (VSPLOT_CRTE), and the number of positions off for all trials (VSPLOT_OFF) was measured for each subject.

### 2.2. Preprocessing and feature extraction

We preprocessed the rs-fMRI with a standard pipeline and extracted dFNC features from the preprocessed data.

#### 2.2.1. Data processing

For processing the fMRI data, statistical parametric mapping (SPM12, http://www.fil.ion.ucl.ac.uk/spm/) was used. In this process, we first implemented a slice-timing correction on the fMRI data. We then applied rigid body motion correction using an SPM toolbox to correct subject head motion. In the next step, we performed spatial normalization to echo-planar imaging (EPI) template in the standard Montreal Neurological Institute (MNI) space and resampled it to 3×3×3 mm. Finally, we used a Gaussian kernel with a full width at half maximum (FWHM) = 6 mm to smooth the fMRI images.

To extract reliable and fully automated intrinsic connectivity networks (ICNs), we used the Neuromark templates as input to a spatially constrained ICA approach [15]. We extracted 53 ICNs and then grouped them into seven domains based on preexisting anatomical and functional information. These domains include the subcortical network (SCN), auditory network (ADN), sensorimotor network (SMN), visual network (VSN), cognitive control network (CCN), default-mode network (DMN), and cerebellar network (CBN). The 53 extracted ICNs were further described in [16].

#### 2.2.2. Dynamic functional network connectivity (dFNC)

For each subject, *i = 1 … N*, whole-brain dFNC was estimated via a sliding window approach, as shown in Fig.1 [17]. We used a tapered window, which was obtained by convolving a rectangle (window size = 20TRs = 14.4 s) with a Gaussian (σ = 3 s), to localize the dataset at each time point. The covariance matrix among the 53 components was calculated to measure the FNC between ICNs for each window (Fig. 1 Step1). We calculated 1378 connectivity features within each window. Next, we concatenated dFNC estimates of each window for each subject to form a *C × C × T* array (where *C=53* denotes the number of ICNs, and T=1019 indicates the number of windows). The array represented the changes in brain connectivity between ICNs as a function of time (Fig.1 Step 1).

**Fig. 1.**
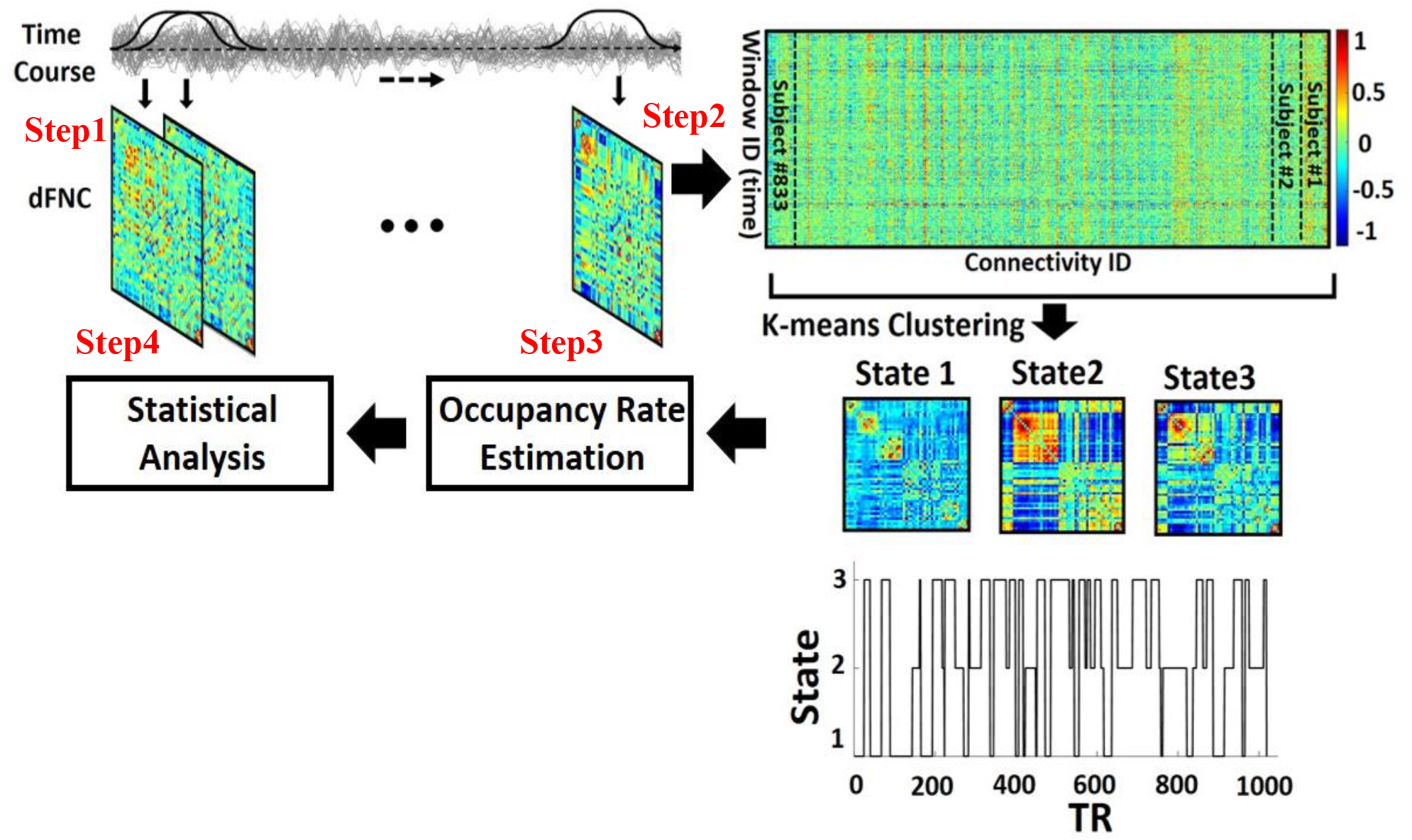
Analytic pipeline used in this study. Step 1 indicates the process by which connectivity matrices were extracted from overlapping windows. Within each window, the Pearson correlation among all components was calculated. These connectivity matrices were concatenated (Step 2), and k-means clustering was applied. States and a state vector were identified over time. The state vector showed how the functional network connectivity, or the state of the brain network, changed over the time. Lastly, the occupancy rate for each state was calculated on a per-subject basis, and statistical analyses were performed (Step 3). Partial correlation, in which we accounted for age and sex effects, was used to find a link between dynamic functional network connectivity features and spatial orientation test scores (Step 4).

### 2.3. Data analysis

A clustering method was applied to the dFNC values to identify dynamic states. The portion of time that each subject spent in each state was calculated, and statistics were performed to analyze the relationship between state occupancy and spatial orientation skills.

#### 2.3.1. Clustering and occupancy rate feature estimation

We next concatenated the dFNC of all subjects, as shown in Step 2 of Fig.1, and applied the *k*-means clustering algorithm to the dFNC windows to partition the data into sets of distinct clusters representing transient connectivity “states” [18,19]. The optimal number of centroid states was estimated using the elbow criterion based on the ratio of within to between cluster distances. By sweeping the k-value from 2 to 9, we found that the optimal number of clusters was 3. The dFNC correlation was also used as a distance metric in this k-means clustering algorithm with 100 iterations. After separating each of the dFNC samples into states, we calculated the state-specific occupancy rate (OCR) for each subject. The OCR represents the portion of time that each subject spent in a given state (Step 3 in Fig.1). In addition, we averaged the dFNC features of each subject in each state. It is worth mentioning that each subject had multiple dFNC samples in each state, which enabled us to average all of the dFNC features (i.e., the average of 1378 connectivity features) of each subject to obtain their state-specific FNC.

#### 2.3.1. Statistical analysis

To find a link between OCR features and the spatial orientation scores, we used partial correlation accounting for age and sex (Step 4 in Fig.1). We performed statistical analyses on all 3 OCR features and used the Benjamini-Hochberg correction method for false discovery rate (FDR) correction of all p-values [20]. We additionally used partial correlation accounting for age and sex to determine whether there was a relationship between the spatial orientation scores and state-specific FNC. We applied FDR correction to all p-values [20].

## RESULTS

This section describes the results for analyzing age-related differences in spatial orientation and identifying distinctive neural states and their link with spatial orientation scores.

### 3.1. The relationships of age with spatial orientation

Results showed that variation in age corresponded to variation in spatial orientation. The number of accurate alignments (VSPLOT_TC) decreased with age (R=-0.097, p=0.005). The number of positions away from alignment (VSPLOT_OFF, R=0.072, p=0.037) and the ratio of the median reaction time with the expected number of clicks to correctly align the lines (VSPLOT_CRTE, R=0.098, p=0.004) both increased with age.

### 3.2. Overview of dFNC states identified by k-means

We identified 3 distinct dFNC states (Fig. 2) that occurred across all subjects. State 1 occurred most frequently, followed by state 3 and state 2. Each of the 3 states incorporates highly distinctive activation patterns. In general, state 1 was characterized by relatively weak connectivity levels both between and within regions, with the exception of low to moderate levels of positive dFNC within the SCN, VSN, and SMN. State 2 displayed higher levels of positive functional connectivity within-SCN, within-CBN, and inter-SCN-CBN than those of states 1 and 3. Additionally, state 2 had high levels of positive and negative connectivity between most regions and the later CCN components. Lastly, state 3 showed within-VSN, inter-VSN-SMN, within-DMN, within-ADN, inter-ADN-VSN, and inter-ADN-SMN levels of positive connectivity that exceeded those of states 1 and 2. Both states 2 and 3 were characterized by high levels of negative functional connectivity inter-SMN-CBN, inter-CBN-VSN, inter-SMN-DMN, and inter-VSN-DMN. However, state 3 showed inter-DMN-ADN, inter-DMN-SMN, and inter-DMN-VSN negative connectivity that was much more negative than states 1 and 2. Additionally, the SCN showed connectivity levels with other networks for state 3 that were generally between state 1 and state 2.

**Fig. 2.**
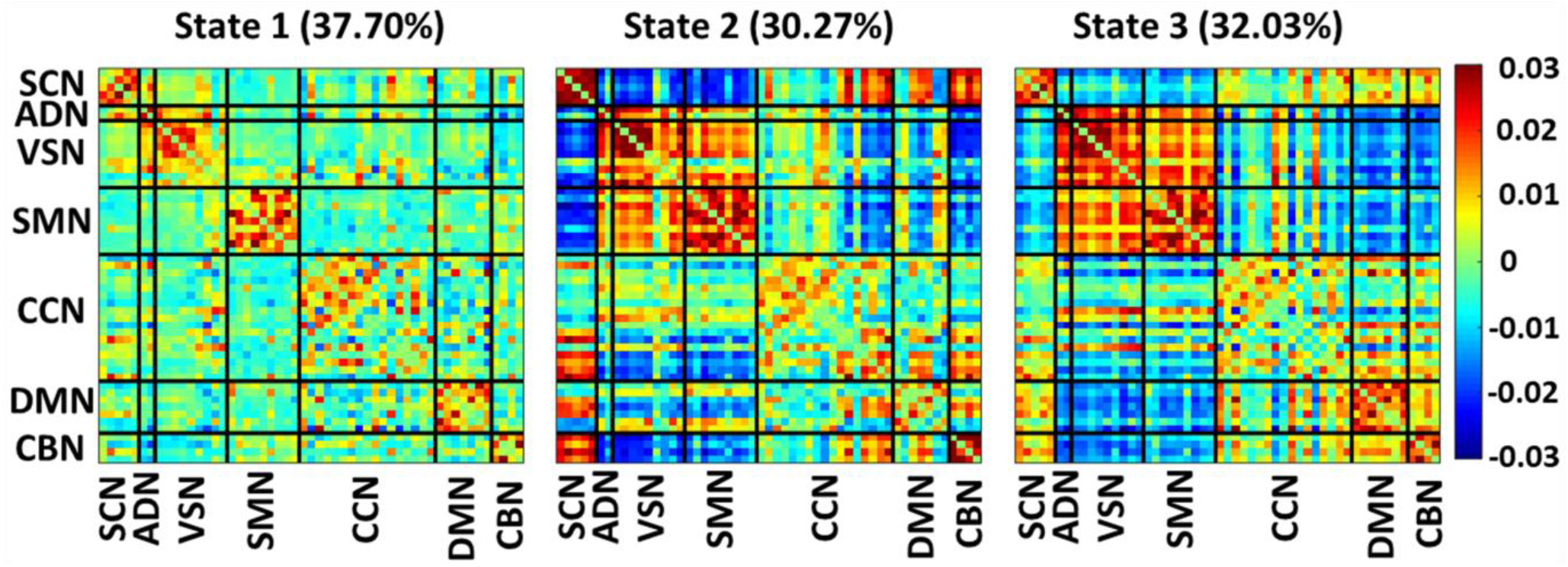
dFNC States identified by k-means. The above heat maps indicate the dFNC within and between brain networks for each of the 3 identified states. The percentage to the right of each state number indicates the portion of the recordings across all subjects and time windows belonging to a particular state. The labels on the x- and y-axes indicate the networks from which dFNC was extracted. The colorbar on the right side of the figure corresponds to the level of connectivity shown in the heat maps. The color bar shows the strength of the connectivity. SCN: Subcortical network, ADN: auditory network, VSN: visual network, SMN: sensorimotor network, CCN: cognitive control network, DMN: default-mode network, and CBN: cerebellar network.

### 3.3. The link between spatial orientation scores and OCR

The occupancy rate associated with a subject’s time in state 3 demonstrated a statistically significant level of correlation with the VSPLOT_TC (Table 1). In more detail, we found that staying in state 3 has a positive link with the total number of the correct trials in the spatial orientation test (r=0.103, FDR p=0.009). However, neither the occupancy rate of state 1 nor that of state 2 had significant relationships with the spatial orientation scores. The correlation between the occupancy rate of a subject for state 3 and VSPLOT_OFF was closer to significant than any of the other remaining states and spatial orientation scores but was still not significant after FDR correction.

**Table 1.** Correlation between OCR and spatial orientation scores

|  | State1 | State2 | State3 |
| --- | --- | --- | --- |
| **VSPLIT_TC (R)** | -0.048 | -0.021 | **0.103** |
| **VSPLIT_TC (p)** | 0.249 | 0.538 | **0.009** |
| **VSPLIT_CRTE (R)** | -0.012 | 0.007 | 0.006 |
| **VSPLIT_CRTE (p)** | 0.865 | 0.865 | 0.865 |
| **VSPLIT_OFF (R)** | 0.034 | 0.013 | -0.071 |
| **VSPLIT_OFF (p)** | 0.475 | 0.692 | 0.114 |

**Note:** OCR: occupancy rate, VSPLIT: variable short Penn line orientation test, VSPLIT\_TC: The total number of the correct trials, VSPLIT\_CRTE: median reaction time divided by the expected number of clicks to correctly align the lines, VSPLIT\_OFF: number of positions off for all trials. R: Pearson correlation, p: p-value, bold face numbers are the significant statistics after FDR correction.

### 3.4. The link between spatial orientation scores and cell-wise FNC in each state

As shown in Fig. 3, the analysis of the correlation between the individual cells of each state and VSPLOT_OFF found that a large number of cells across all states had significant levels of correlation with VSPLOT_OFF when not FDR corrected, and multiple cells of the within-SMN FNC for states 1 and 2 were significantly negatively correlated with VSPLOT_OFF after FDR correction. Multiple cells of within-SMN dFNC for state 3 were also significant without FDR correction but not significant following FDR correction. We examined the correlation between the VSPLOT_TC and VSPLOT_CRTE scores and the cell-wise dFNCs as well, and we did not identify any significant relationships there.

**Fig. 3.**
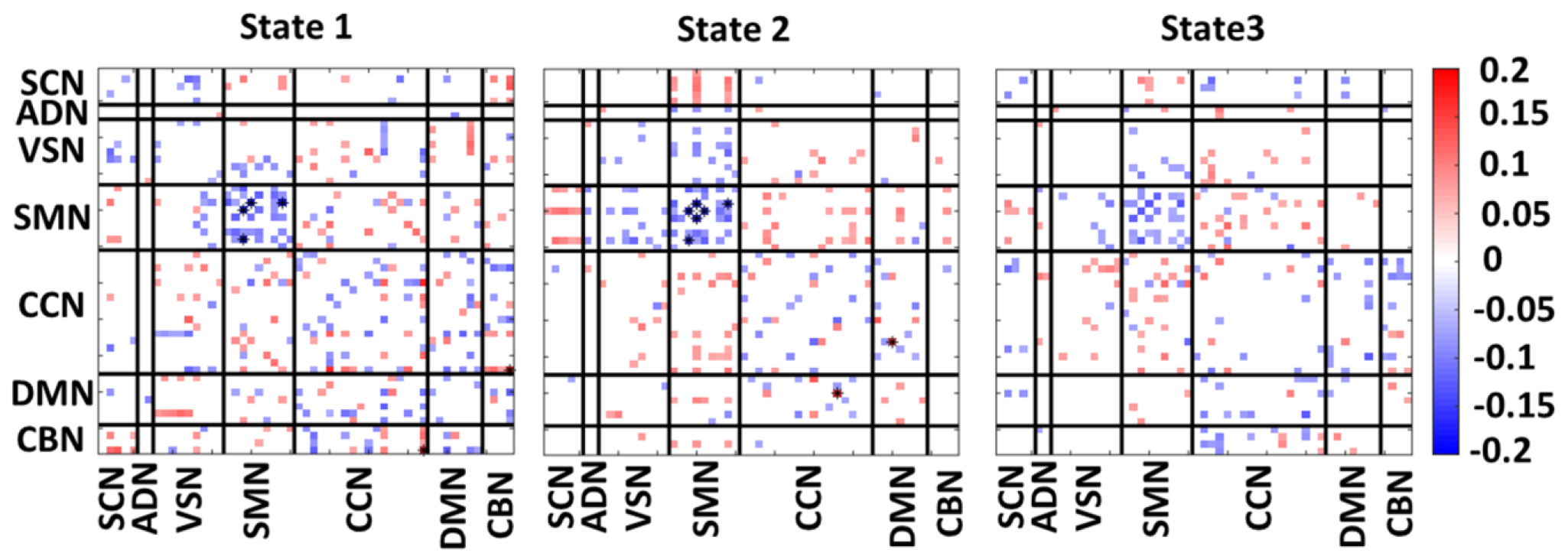
VSPLOT_OFF Correlation with state-specific FNC. The labels on the x-axis and y-axis indicate the domains from which the dFNC was extracted. The colormap on the right indicates the degree of correlation associated with each cell that had a significant correlation without FDR correction. The stars overlapping some of the cells indicate that those cells had a significant level of correlation with the VSPLOT_OFF after FDR correction. The color bar shows the strength of the connectivity difference. SCN: Subcortical network, ADN: auditory network, VSN: visual network, SMN: sensorimotor network, CCN: cognitive control network, DMN: default-mode network, and CBN: cerebellar network.

## DISCUSSION

In this study, we used a data-driven approach and extracted 53 ICs for the whole brain and used a sliding window approach followed by a clustering method to study brain dFNC of subjects within the age range of 22 years to 37 years. We identified three distinct dFNC neural states and found that the subjects’ brains in this age range were highly dynamic. We found that functional connectivity within networks, such as the SCN, CCN, DMN, and CBN, was more active than the between-network connectivity. Interestingly, the functional connectivity between sensory networks including the ADN, VSN, and SMN showed more temporal change than other within-network connectivity. Additional studies are needed to compare the dynamical properties of the brains of healthy subjects across a greater age range.

We further investigated the link between spatial orientation scores with age and also with dFNC properties. We found that performance on spatial orientation tests decreased with age across three metrics, which is well-established in existing literature [6]. However, previous literature did not study the effect of age on spatial orientation scores of subjects in the current study’s age range. Because of these relationships within our dataset, we controlled for age in the statistical analyses comparing dFNC states to spatial orientation scores.

The amount of time that subjects spent in state 3, with higher connectivity in sensory networks, significantly correlated with their total number of correct VSPLOT trials. The spatial orientation task is likely to integrate the information of multisensory signals, including visual sensory and somatosensory [21]. It is also noteworthy that high levels of SMN, VSN, and ADN connectivity characterize state 3, given that those networks are associated with proprioceptive and vestibular systems that are known to play an important role in spatial orientation [22]. Additionally, we found that multiple cells within the SMN of states 1 and 2 had significant levels of negative correlation with VSPLOT_OFF. This finding, combined with our other results that showed that the state with the highest SMN functional connectivity was positively correlated with spatial orientation, provides strong evidence that an increase in functional connectivity within the SMN is associated with improved spatial orientation.

Also, it was noteworthy that state 3 had higher levels of within-DMN connectivity than the other states. The DMN includes the posterior cingulate cortex that has been associated with aspects of spatial orientation and of which some researchers have claimed the retrosplenial cortex should be considered a subregion [23]. The retrosplenial cortex has been associated with the formation and use of cognitive maps [24–26], which play an integral role in spatial navigation. Additionally, state 3 is characterized by relatively higher functional connectivity between DMN and CCN. Further investigation is needed to explore the role of DMN/CCN connectivity in spatial orientation.

There are a few limitations to this study. The choice of window size involves an implicit assumption about the dynamical behavior of the networks in that a short window captures more rapid fluctuations, whereas a longer window performs more smoothing relative to a shorter one. Future work can be conducted to evaluate the range of dynamics more comprehensively [27]. In addition, k-means clustering requires that some of the clustering parameters, including distance metrics, be predefined. When applying different distance metrics, we did not observe a significant difference in the results; however, using other clustering methods such as robust continuous clustering would eliminate this shortcoming and potentially improve the clustering results [28]. Other future directions for this study include (1) examining the reproducibility of the identified states across multiple recordings with the same subjects, (2) looking at whether similar states exist in different age groups, and (3) examining whether those states occur at the same rate in other age groups.

## CONCLUSIONS

Previous literature studied static functional connectivity and its association with spatial orientation performance. In the work reported here, we extend this existing body of knowledge into the realm of dynamics, investigating how time-varying properties of the whole-brain functional network connectivity relate to spatial orientation performance in young healthy subjects. We identified and characterized 3 distinct dFNC states across healthy young adults. We found that the amount of time that a subject spent in one of those states, a state with higher sensory network connectivity, correlated with improved performance on the variable short Penn line orientation test, a test commonly used to assess spatial orientation capabilities. The identified state had increased within-DMN FNC, which warrants further investigation. Additionally, we found a significant correlation between SMN FNC values and performance on the variable short Pen line orientation test. These findings provide new insights into how spatial orientation is related to brain activity on a network level over time.

## COMPLIANCE WITH ETHICAL STANDARDS

This study was approved by the institutional review board of the Washington University - University of Minnesota Consortium of the Human Connectome Project (WU-Minn HCP).

## ACKNOWLEDGMENTS

We thank the Washington University - University of Minnesota Consortium of the Human Connectome Project (WU-Minn HCP) for collecting the data.

